# Joint and tissue mechanics in post-traumatic osteoarthritis: insights from the rat model

**DOI:** 10.1101/2024.06.27.601039

**Authors:** Judith Piet, Amir Esrafilian, Seyed Ali Elahi, Falk Mielke, Maarten Van Nuffel, Gustavo Orozco, Sanne Vancleef, Rik Lories, Rami K. Korhonen, Peter Aerts, Sam Van Wassenbergh, Ilse Jonkers

## Abstract

**Objective:** Altered mechanical loading is a known risk factor for osteoarthritis. Destabilization of the medial meniscus (DMM) is a preclinical gold standard model for post-traumatic osteoarthritis and is thought to induce instability and locally increased loading. However, the joint- and tissue-level mechanical environment underlying cartilage degeneration remains poorly documented.

**Design:** Using a custom multiscale modeling approach, we assessed joint and tissue biomechanics in rats undergoing sham surgery and DMM. High-fidelity experimental gait data were collected in a setup combining biplanar fluoroscopy and a ground reaction force plate. Knee poses and joint-level loading were estimated through musculoskeletal modeling, using bony landmarks, semi-automatically tracked via deep learning on fluoroscopic images, and ground reaction forces. A musculoskeletal model of the rat hindlimb was adapted to represent knee flexion-extension, valgus-varus, and internal-external rotation. The tissue-level cartilage mechanical environment was then spatially estimated, using the musculoskeletal modeling parameters as inputs into a dedicated finite element (FE) model of the rat knee, comprising cartilage and meniscal tissues. Experimental gait data and modeling workflows, including musculoskeletal models and FE meshes, are openly shared through a data repository.

**Results:** In rats with DMM, the frontal plane knee pose was altered, yet there was no indication of joint-level overloading. Tissue-level mechanical cues typically linked with cartilage degeneration were not increased in the medial tibial cartilage, despite evidence of tissue structural changes.

**Conclusion:** DMM did not increase joint and tissue mechanical responses in the knee medial compartment, suggesting that mechanical loading alone does not explain the observed osteoarthritis-like structural changes.

## Introduction

Osteoarthritis is a complex multi-factorial disease that affects various joint tissues, including articular cartilage, subchondral bone, menisci, and synovium ^1^. The disruption of the mechanical and biological joint environment contributes to cartilage degeneration, a hallmark of osteoarthritis^2^. Inadequate or excessive mechanical loading are widely accepted, but still poorly understood, drivers of the onset and progression of cartilage damage during osteoarthritis. Articular cartilage is a mechano-adaptive tissue, and *in vivo* and *in vitro* experiments have shown that its anabolic and catabolic activity is regulated by mechanical loading (^3,4^, review in ^5^). Furthermore, the molecular mechano-regulatory pathways within cartilage are disrupted in osteoarthritis (^6^, review in ^7^).

The cartilage mechanical environment is complex, encompassing both a solid matrix and a fluid phase. Various mechanical cues, such as excessive concentrated loads, excessive fluid exudation, and excessive compressive, shear, and collagen fibril deformation, are hypothesized to contribute to local cartilage degeneration by causing loss of fixed charge density from the proteoglycan matrix molecules and disorganization of the collagen fiber matrix ^8–12^.

To study cartilage changes in post-traumatic osteoarthritis, the current preclinical gold standard is the destabilization of the medial meniscus (DMM) in the rodent knee, which triggers progressive joint degeneration ^13–15^. DMM is typically referred to as a model of joint instability with increased mechanical loading leading towards osteoarthritis in both mice ^13,16^ and rats ^14,17^.

Previous studies have tried to understand the complex mechanical environment in the knee joint and cartilage tissue of preclinical models. These used advanced biomechanical modeling techniques like musculoskeletal and finite element (FE) models, both with and without post-traumatic osteoarthritis. FE models have been developed to analyze cartilage deformations (strains) and concentrated loads (stresses). However, these FE models used simplified input data that did not accurately represent real movement, as high-fidelity species-specific movement data are scarce ^18–21^. Musculoskeletal models have been used to describe movement based on joint positions and ground reaction forces (i.e., contact forces between the paw and ground), but relied on marker-based motion capture to describe joint positions ^22^. The accuracy of this technique is limited because it assumes that skin markers move with the underlying skeleton, which is contradicted by the presence of significant soft tissue artifacts. Tracking skeletal landmarks directly on fluoroscopic images gives more accurate knee angles, but this requires a dedicated and complex setup ^23^.

Therefore, a comprehensive and accurate description of the multiscale joint- and tissue-level mechanical environment in rats following DMM is still lacking, limiting our understanding of the role of mechanical loading on *in vivo* cartilage degeneration in post-traumatic osteoarthritis. To address this research gap, we characterized the spatial distribution of joint- and tissue-level mechanics during gait in rats following DMM using high-fidelity, species-specific data on joint pose and ground reaction forces.

## Method

All experiments with rats were approved by the Ethics Committee for Animal Research (KU Leuven and Universiteit Antwerpen, Belgium, P134/2018). Twelve male Sprague-Dawley rats were purchased from Envigo at 11 weeks of age and experienced a one-week acclimatization period, with brief daily handling by the main researcher. They were housed by two in individually ventilated cages at KU Leuven, and fed ssniff^®^-R/M-H.

### DMM and implantation of radio-opaque markers

At 12 weeks of age (384±12 g body weight), rats were randomly distributed into two groups of 6 rats (simple randomization), to undergo DMM or sham surgery on the right knee ^13,14^ (see **Fig. 1A**). DMM surgeries were done under the supervision of a skilled micro-surgeon using magnifying surgical loupes. Two weeks following the knee surgery, at 14 weeks of age, radio-opaque beads were implanted onto the right tibia and femur to ease the tracking of the femur and tibia on fluoroscopic images ^23^ (see **Fig. 1B)**. Surgical procedures are detailed in **Supplementary Text**.

**Fig 1.**
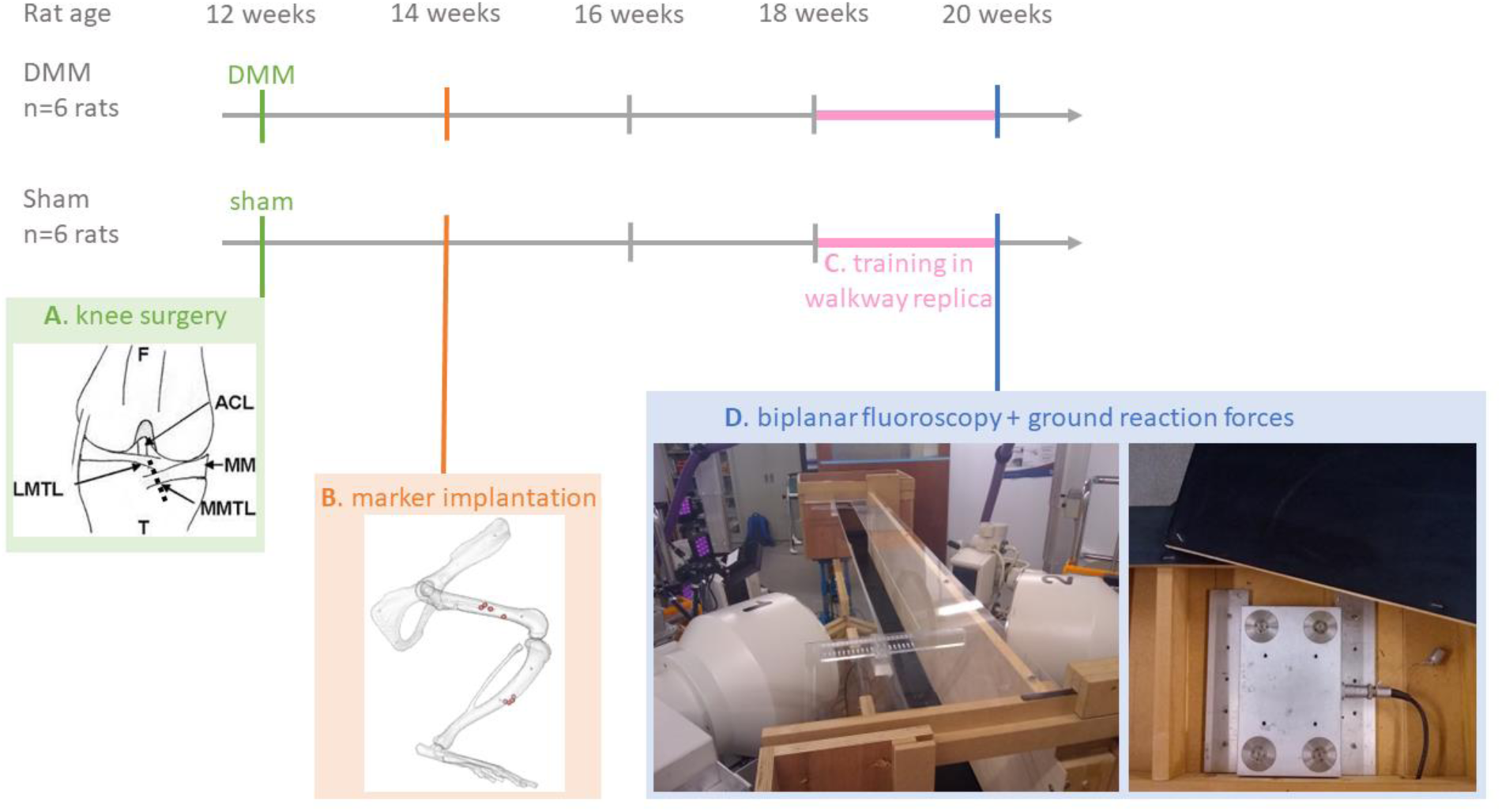
Experimental timeline and groups. (**A**) Rats received DMM or sham surgery on their right knee, (**B**) followed by implantation of radio-opaque beads onto their right femur and tibia. (**C**) After training to walk in a straight line in a walkway, (**D**) their gait was recorded in a setup integrating biplanar fluoroscopy and a force plate.

### Behavioral training

To ensure that rats could ambulate in a straight line in the instrumented walkway during the fluoroscopic recordings, they were trained from 18 weeks to 20 weeks of age to ambulate in a walkway replica (**Fig. 1C**). The rats were trained in a blinded manner to cross the walkway with a consistent gait (trotting without galloping), 5 days per week, for 10-15 minutes a day, using small pieces of Cheerios as a food reward ^24^.

### Ground reaction forces and biplanar fluoroscopy acquisition

To accurately record rat gait, we created a unique experimental platform (3D2YMOX or 3-Dimensional DYnamic MOrphology using X-rays, University of Antwerp ^25^) that simultaneously records calibrated biplanar fluoroscopic video images and single hind limb ground reaction forces using a piezo force plate embedded in the walkway (**Fig. 2**). Using this setup, rat gait was captured during free ambulation, eight weeks following knee surgery (at 20 weeks of age, 461 +/- 19 g) (**Fig. 1D**).

**Fig 2.**
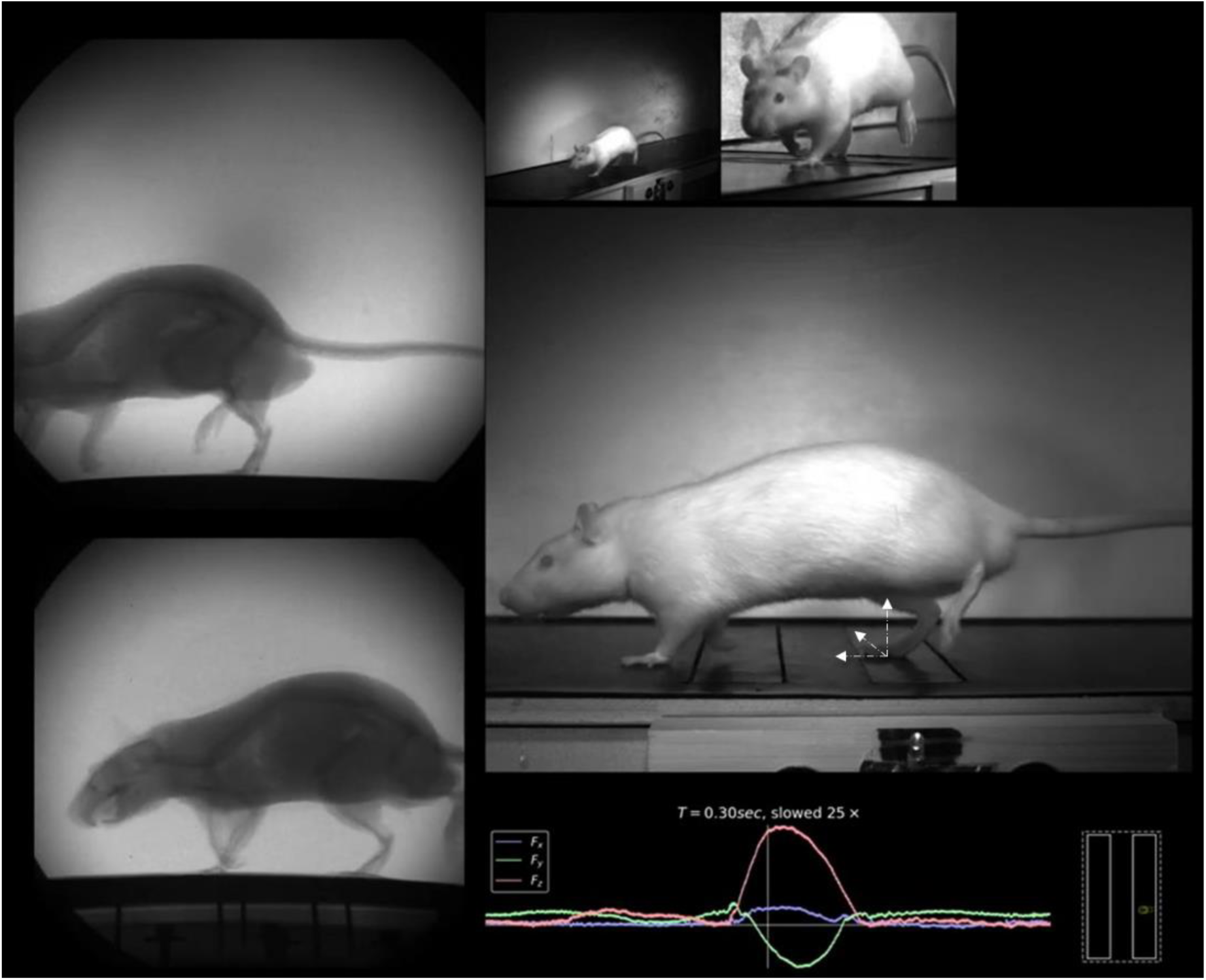
Experimental setup for simultaneous acquisition of biplanar fluoroscopy and ground reaction forces. To the left, synchronous biplanar X-ray images (X-ray sources: 65 and 71kV, 40mA, record rate 500 fps). To increase the chances of only the right foot hitting the force plate, two contact sockets were mounted above the force plate (Kistler “Squirrel” force plate, acquisition rate of 20000 Hz). To the top right, images from high-speed cameras to identify single hindlimb contact with one of the two rectangular force plate sockets. Bottom right, force plate trace and center of pressure location viewed from above.

### Data processing

Rat speed varied throughout the trials, given that over-ground walking data at a self-selected speed were collected . Only trials for which the rats had right foot force plate contact, for which the skeleton of the right hindlimb was in the field of view of the biplanar X-Rays, and for which rats had a consistent trotting pattern (no jumping, slowing down, or pausing on the force plate) were analyzed. In total n=27 gait trials for N=12 rats were analyzed, with inclusion criteria defined prior to analysis and no trials were excluded for other reasons.

Raw force plate signals were filtered with a zero-phase band-pass filter (0.2 to 100 Hz) to remove noise, voltage drift, and mechanical oscillations. Forces, torques, and center of pressure were then computed. The data baseline was identified for each trial and subtracted.

Biplanar X-ray images were processed in XMALab ^26^. The radio-opaque beads surgically implanted on the femur and tibia shafts were tracked manually on all frames, and in a blinded manner in XMALab, with a retro-projection error below 2 pixels or 0.226 mm (see **Fig. 3A**).

**Fig 3.**
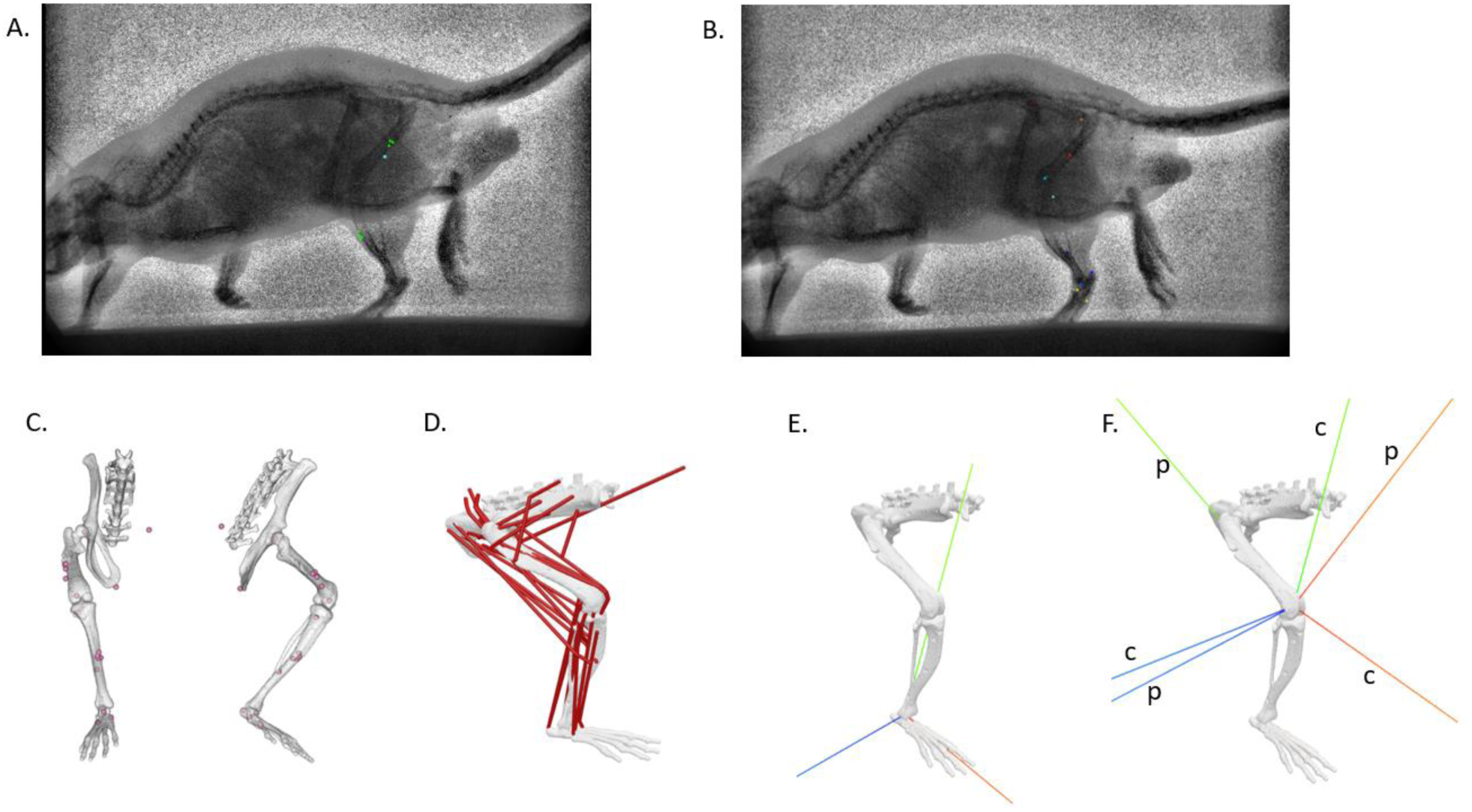
Musculoskeletal workflow. (**A**) Manual labeling of the clusters of implanted bone markers. Images were calibrated using calibration cube images and undistorted using distortion grid images. Image contrast was improved using the CLAHE plugin in FIJI ^43,44^.; (**B**) Automatic labeling of anatomical skeletal landmarks in DeepLabCut (pubic symphysis, left and right femur acetabula, center of right femur condyles, right distal and proximal tibia-fibula junctions, right ankle, right heel, proximal ends of the 3^rd^ and 5^th^ right metatarsals); (**C**) Position of virtual markers (implanted bone markers and anatomical skeletal landmarks) in OpenSim. The animal-specific femur and tibia micro-CT scans were registered to the stereolithography (STL) files from the generic model to identify the position of the implanted bone markers in the musculoskeletal model using Mimics (Materialise, Leuven, Belgium). (**D**) Musculoskeletal model of the rat hindlimb. The model was scaled based on bone measurements in micro-CT scans relative to the generic model and based on the mass of each rat (**E** ) The tibial coordinate system was defined similarly to the generic model (based on bone anatomy). In this study, all musculoskeletal results are presented in this tibial coordinate system. (**F**) updated knee definition (to be consistent with the FE model’s knee coordinate system). c: child frame; p: parent frame.

10 anatomical skeletal landmarks were labeled manually on 427 frames (equivalent to 6 gait trials). The skeletal landmarks on the remaining frames were then labeled using deep learning, with a validated workflow integrating XMALab and DeepLabCut ^27–29^. A network was trained for each camera, using a training fraction of 0.75 and resnet_101. Landmarks labeled by the networks were inspected and corrected manually if needed, and the networks were trained again on this augmented dataset (see **Fig. 3B**). Gait speed of individual trials was calculated using the horizontal displacement of the pubic symphysis skeletal landmark over time.

### Sample imaging and histology

Following gait trials, rats were sacrificed and hindlimbs were analyzed *ex vivo*. To identify the location of surgically implanted beads onto the femur and tibia, dissected operated hindlimbs were imaged with a coarse resolution in a micro-CT scanner (GE Nanotom M, 40 micrometer voxel size, 90 kV, 200 μA).

Knee soft and hard tissues were then imaged with a fine resolution using Hexabrix, which is negatively charged and used as a contrast agent for cartilage ^30,31^ (GE Nanotom M, 6.5 micrometer voxel size, 60 kV, 350 μA).

Tissue damage was then assessed with histology, using hematoxylin-safranin-O staining performed on 10 micrometers thick frontal sections, located in the middle of the joint ^32,33^.

### Musculoskeletal modeling

By combining a musculoskeletal model of the rat hindlimb with our experimental data, we estimated knee contact forces to examine whether DMM resulted in abnormal joint-level loads, and the associated knee poses to examine whether DMM resulted in joint instability.

An OpenSim musculoskeletal model of the rat hindlimb was adapted in OpenSim 4.4 to represent 3 rotational degrees of freedom at the knee (i.e., flexion-extension, valgus-varus, and internal-external rotation), and no translational degrees of freedom ^34–36^ (**Fig 3**, see details in Supplementary Text).

To obtain knee poses and joint-level loading, we estimated knee angles, external joint moments, muscle forces, and knee contact forces during the stance phase (respectively inverse kinematics, inverse dynamics, static optimization of muscle forces distribution, and joint reaction analysis). To estimate loads on the medial tibia, the force on the medial compartment was calculated using the axial contact force, the adduction contact moment, and the intercondylar distance ^37^. To estimate the accumulated loading on the medial compartment during the stance phase, the impulse on the medial compartment was calculated as the integral of force on the medial compartment over the stance phase. Inputs and outputs of the musculoskeletal modeling workflow were validated via comparison with relevant findings from the literature (ground reaction forces, knee angles, knee moments, peak muscle forces, and knee contact forces, see details in **Supplementary Text**).

### FE Analysis

To estimate spatial cartilage mechanical loads, an FE model of the rat knee joint was created in Abaqus (Dassault Systèmes, United States, version 2021, implicit analysis), after manually segmenting the fine-resolution hexabrix scan of a knee from the sham surgery group (**Fig. 4A**), post-processing, and meshing knee structures. Cartilages, menisci, and ligaments were included in the model (**Fig. 4B**). Cartilage was modeled as a fibril-reinforced poro-viscoelastic material with a fluid phase representing the interstitial fluid and a solid phase. The latter was modeled with a hyperelastic nonfibrillar matrix representing the proteoglycans and a viscoelastic fibril network representing collagen fibrils. Menisci were modeled similarly, except that the fibril network was linear elastic (fibril-reinforced poroelastic material). To use representative input parameters to drive the FE model, we selected for each condition (sham surgery or DMM) the gait trial for which the peak contact force magnitude was closest to the group estimated mean. Inputs to the FE models were knee flexion angles, knee contact forces, and adduction moments (generated by muscles, inertial, and gravitational forces) (**Fig. 4C**), obtained from musculoskeletal analysis of the corresponding gait trial (see more details on the FE model in **Supplementary Text**). Additionally, DMM surgery was simulated by deactivating the anterior attachment of the medial meniscus horn.

**Fig 4.**
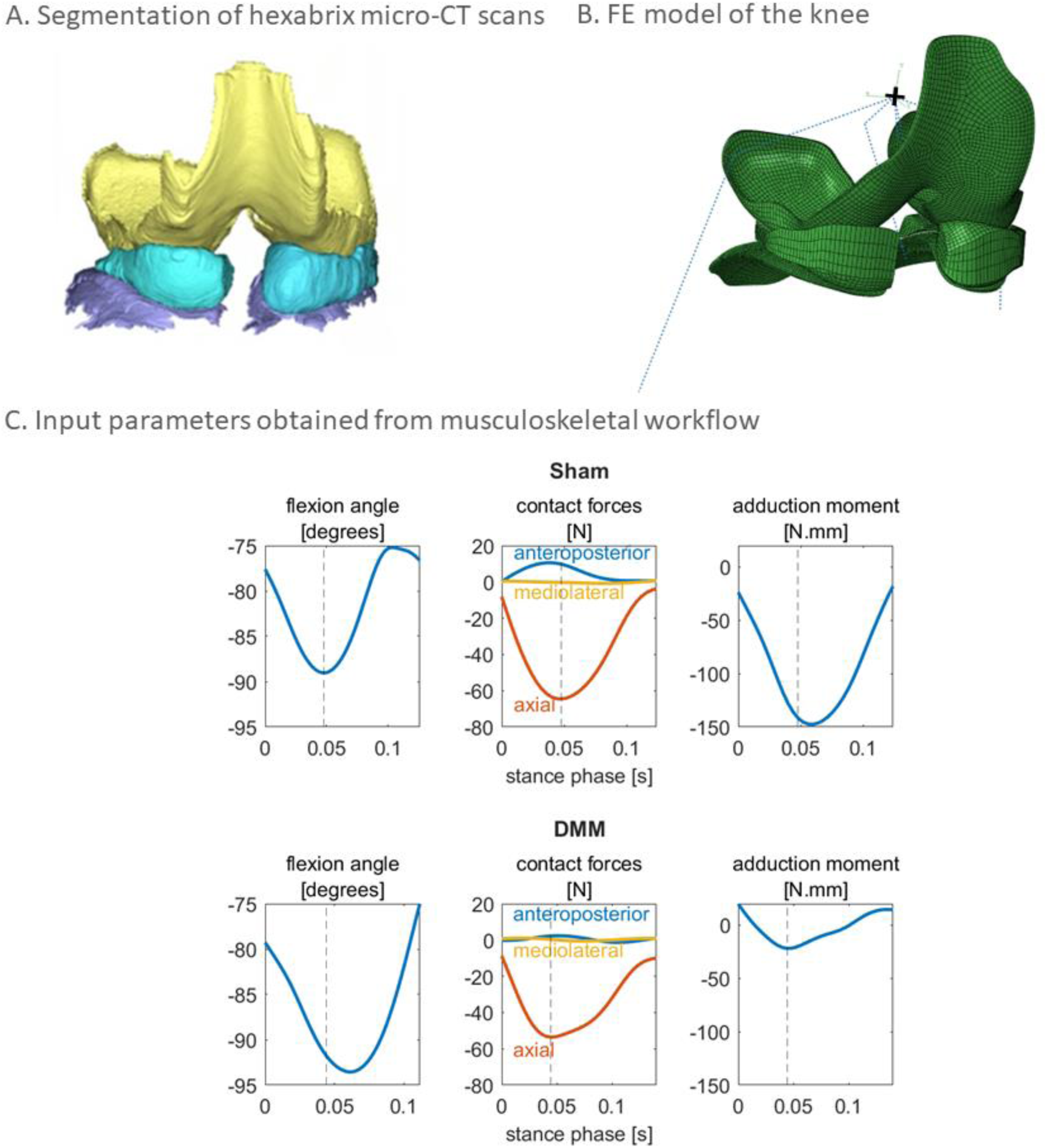
FE model. **(A)** Knee femoral cartilage (yellow), tibial cartilage (purple), and menisci (blue) were segmented on fine-resolution hexabrix-enhanced micro-CT images. **(B)** Structures were meshed using hexahedral elements (C3D8P, femoral cartilage = 12,318 elements, menisci = 6,248 elements, tibial cartilage = 21,555 elements). Knee ligaments and reserve stiffnesses were modeled using connectors (i.e., nonlinear elastic) ^45^. The reference point of the model is represented with the black “+” sign. **(C)** Bottom nodes of the tibial cartilage were fixed, while knee flexion angle, contact forces and resultant knee abduction/adduction moment were applied to the femoral cartilage reference point. Outputs of the FE analyses were analyzed when the magnitude of the knee contact force was at its maximum (vertical dashed line).

FE outputs were examined at the peak magnitude of the knee contact force on the medial compartment. To estimate the location of loading, the center of pressure in the medial tibia cartilage was calculated using contact pressures between the femur cartilage and medial meniscus, and the medial tibial cartilage ^38^. The distribution and magnitude of specific mechanical cues associated with cartilage degeneration were assessed: fluid velocities, maximum shear and compressive strains, maximum principal stress, and strains in the direction of collagen fibrils ^8–11^.

### Statistical Analysis

#### Gait data and musculoskeletal analysis

Forces and moments were normalized to each rat’s body weight (BW). For each trial, we identified the peak magnitude of the ground reaction force. We also identified the peak magnitude of the knee contact force. We then identified the corresponding knee pose and other contact forces (when the magnitude of the knee contact force is at its maximum). We used R and *lme4* ^39,40^ to perform a linear mixed effects analysis of the relationship between variables of interest (speed, knee adduction moment impulse, knee kinematics and normalized contact forces when the magnitude of the knee contact force is at its maximum), and knee surgery (sham or DMM). As the fixed effect, we entered surgery into the linear mixed effects model. As random effects, we had intercepts for each rat. Visual inspection of residual plots did not reveal any obvious deviations from homoscedasticity or normality. P-values were obtained by likelihood ratio tests of the full model with the effect in question against the model without the effect in question ^41^. The effect size was estimated using the coefficient for the fixed effect β and its standard error SE, representing the average difference between the two groups.

#### FE analysis

To compare mechanical cues between sham and DMM cases, node outputs from the medial compartment at peak magnitude of the knee contact force were analyzed. Only the upper half of nodes with the highest loading (i.e., two upper quartiles) were considered to focus on load-bearing regions, disregarding the lower half with minimal loading. A non-parametric two-tailed rank-sum test assessed differences in median mechanical cues between the two cases, testing the null hypothesis of equal medians, using Matlab (version 9.9.0.1467703, R2020b, Natick, Massachusetts: The MathWorks Inc.). The effect size r was calculated by converting the U statistic from the rank-sum test to a z-value and then dividing by the square root of the total number of observations, providing a standardized measure of the effect size. The probability that a randomly selected data point in group 1 (i.e., sham), is larger than a randomly selected data point in group 2 (i.e., DMM) was evaluated by calculating the probability of superiority.

## Results

### Rats with DMM develop cartilage and bone damage 8 weeks after surgery

Our results confirmed that the DMM model resulted in osteoarthritis-like changes over a two-month interval. Fine-resolution scans of the rat knees, 8 weeks after DMM, showed evidence of structural cartilage damage on the medial tibial plateau. Rats with DMM showed increased penetration of hexabrix staining into the articular cartilage, which suggests loss of cartilage fixed charge density. The animals also had evidence of subchondral bone changes, such as bone cysts, osteophytes, and perforations(see **Fig. 5A**). Histology confirmed proteoglycan loss and abnormal subchondral bone structure (see **Fig. 5B**).

**Fig 5.**
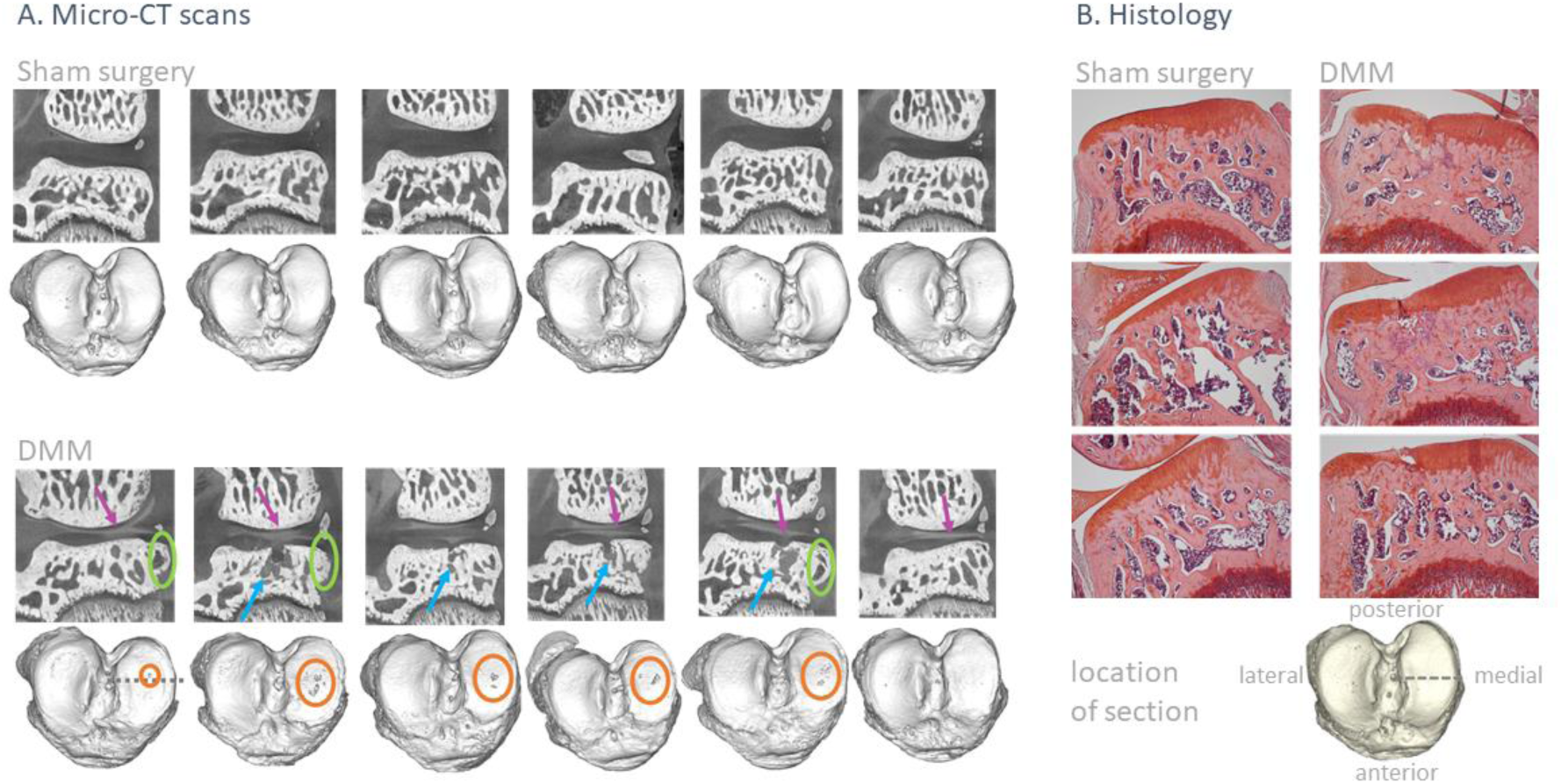
Evidence of osteoarthritis-like structural changes in rats with DMM. **(A)** Micro-CT images of the medial plateau, and 3D reconstructions of the distal tibia. Rats with DMM showed evidence of loss of fixed charge density (purple arrows) in the cartilage of the medial plateau, as well as bone cysts (blue arrows), osteophytes (green circles), and subchondral bone plate perforations (orange circles) in the medial plateau. The dotted grey line on the first 3D reconstruction indicates where images of the medial plateaus were taken. N=6 rats for DMM, N=6 rats for sham. **(B)** Histology section of the medial tibial plateau. Rats with DMM showed evidence of loss of fixed charge density (lighter safranin-O staining) and abnormal bone remodeling. N=3 rats for DMM, N=3 rats for sham.

### Gait speed and limb loading were not different in rats with DMM or sham surgery

Individual traces over the stance phase for ground reaction forces are presented in **Suppl. Fig. S1**. We found that DMM surgery did not significantly affect gait speed and normalized ground reaction force magnitude (i.e., the force that the hind paw exerts on the ground) (**Suppl. Fig. S2**, χ2(1) = 0.04, p = 0.839, β ± SE = -1.1 ± 5.6 cm/s; χ2(1) = 0.02, p= 0.886, β ± SE = 0.0 ± 0.0 BW i.e., Body Weights, respectively).

### DMM did not increase joint loading, but altered joint pose

Individual traces over the stance phase for the outputs of the musculoskeletal workflow are presented in **Suppl. Fig. S3 to S7**.

DMM did not significantly affect the peak magnitude of the resultant knee contact force during the stance phase, χ2(1)= 1.48, p= 0.222(β ± SE = -1.9 ± 1.4 BW). DMM did not significantly affect the axial and mediolateral components of the knee contact force (χ2(1) = 1.48, p =0.223, β ± SE = 1.9 ± 1.4 BW; χ2(1) = 3.12, p =0.077, β ± SE = -0.3 ± 0.2 BW respectively), but it affected the associated anteroposterior component (χ2(1) = 0.22, p = 0.008), lowering it by about 1.1 ± 0.3 BW. DMM did not significantly affect the load distribution between the medial and lateral compartments at peak loading (medial contact force: χ2(1) = 0.07, p =0.789, β ± SE = -0.5 ± 1.9 BW; lateral contact force: χ2(1) = 1.03, p = 0.309, β ± SE = 2.5 ± 2.4 BW respectively). DMM did not significantly affect the loading accumulated on the medial compartment during the stance phase, (medial compartment impulse: χ2(1) = 0.17, p = 0.679, β ± SE = -0.0 ± 0.1 BW.s) (**Fig. 6**).

**Fig 6.**
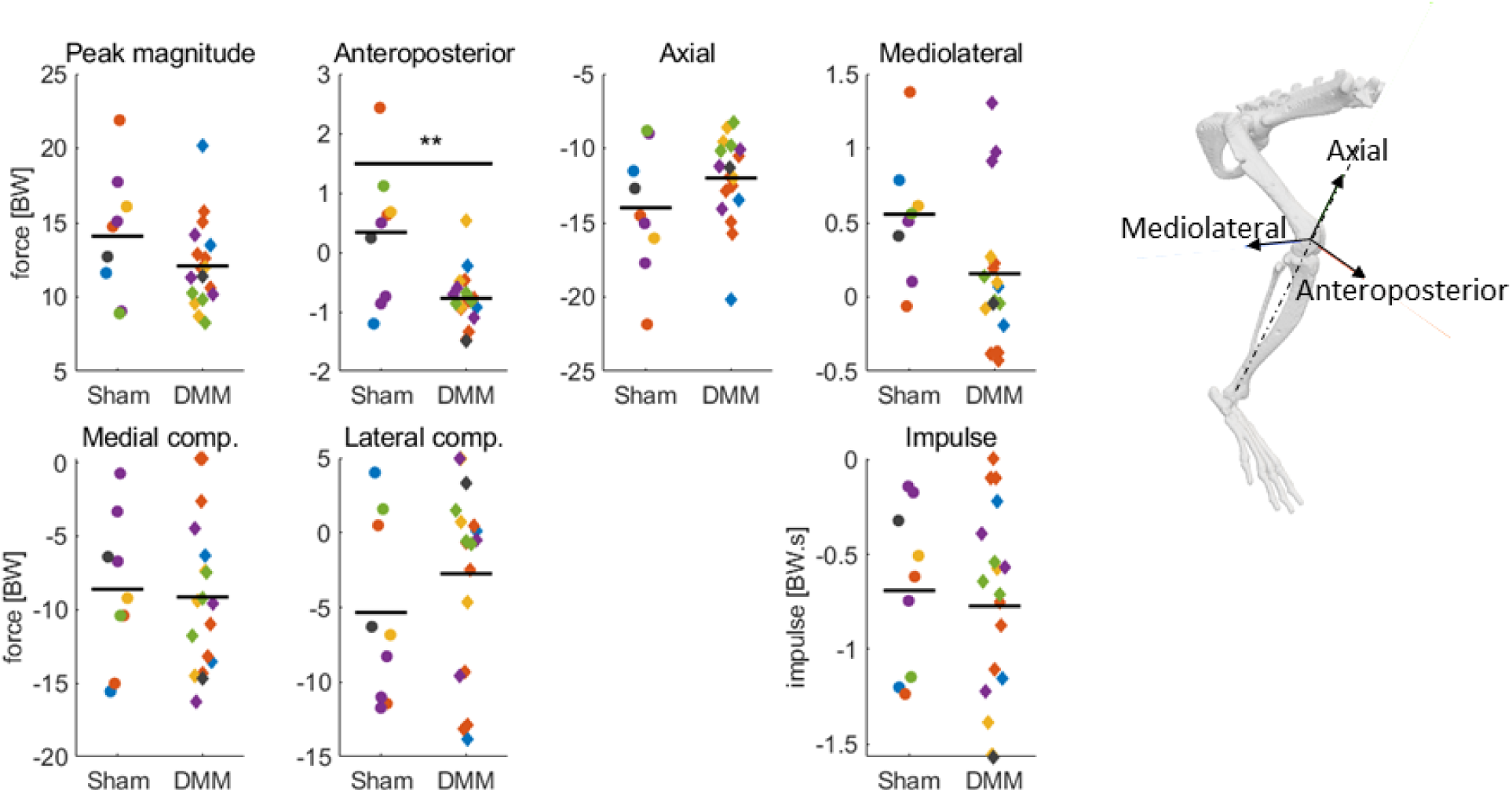
Normalized knee contact forces, and impulse on the medial compartment, at peak magnitude of knee contact force. N=6 rats, n=18 trials for sham; N=6, n=9 for DMM. Trials from the same rat are represented by the same color. The mean for each group is represented with the horizontal black line. Knee contact forces are the forces exerted by the femur on the tibia, expressed in the tibial coordinate system.

Joint instability was confirmed in the frontal plane, as DMM affected the associated adduction angle (χ2(1) = 7.12, p = 0.007), lowering it by about -7.1 ± 2.3 degrees. However, the flexion and internal-external rotation angles were not significantly affected by DMM (χ2(1) = 0.92, p = 0.335, β ± SE = -3.4 ± 3.5 degrees; χ2(1) = 2e-04, p = 0.988, β ± SE = 0.0 ± 1.8 degrees respectively) (**Fig. 7**).

**Fig 7.**
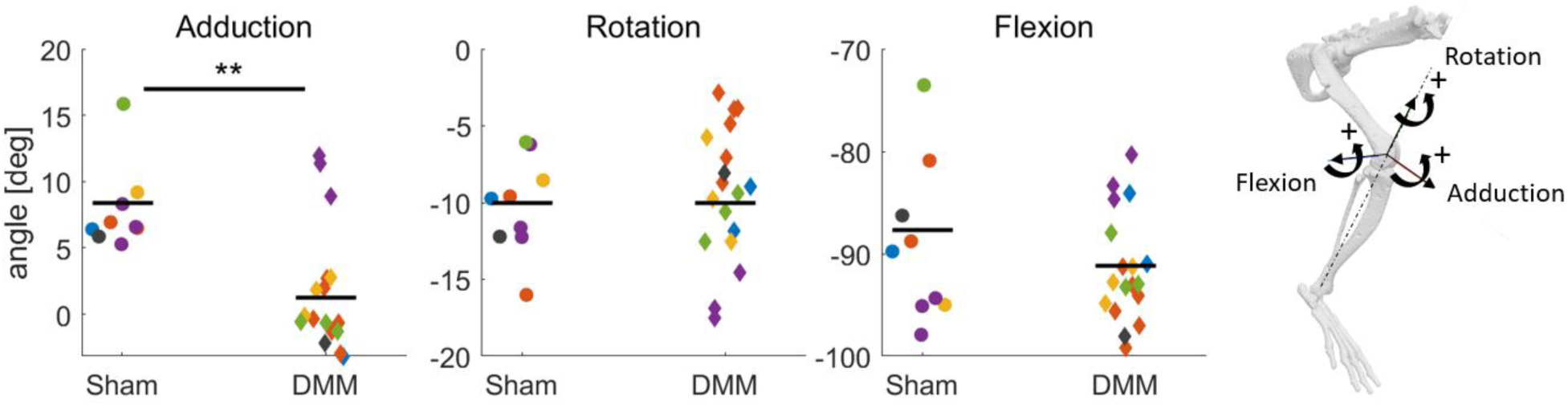
Knee pose at peak magnitude of knee contact force. N=6 rats, n=18 trials for sham; N=6, n=9 for DMM. Trials from the same rat are represented by the same color. The mean for each group is represented with the horizontal black line.

### Mechanical cues typically associated with cartilage degeneration were lower in DMM

Contact pressures between the medial meniscus and the tibial cartilage as well as between the femoral cartilage and the tibial cartilage were higher in sham surgery compared to DMM, but the location of the center of pressure did not change (**Fig. 8B**). With DMM, the examined mechanical cues in the medial tibial plateau were lower compared to sham: fluid velocity (0.0347 +/- 0.0522 mm/s for sham; 0.0249 +/- 0.0445 mm/s for DMM; reporting median +/- median absolute deviation; p<0.0001; effect size r = 0.242; probability of superiority PS = 0.639), maximum principal stress (8.4810 +/- 10.1682 MPa for sham; 6.1073 +/- 7.9730 MPa for DMM; p<0.0001; r = 0.213; PS = 0.623), minimum principal strain (0.0625 +/- 0.0575 for sham; 0.0457 +/- 0.0530 for DMM; p<0.0001; r = 0.213; PS = 0.623), strain in fibril direction (0.0039 +/- 0.0047 for sham; 0.0027 +/- 0.0039 for DMM; p<0.0001; r = 0.171; PS = 0.598), and maximum shear strain ( 0.1175 +/- 0.0863 for sham; 0.0906 +/- 0.0805 for DMM; p<0.0001; r = 0.215; PS = 0.624) (**Fig. 8C**).

**Fig 8.**
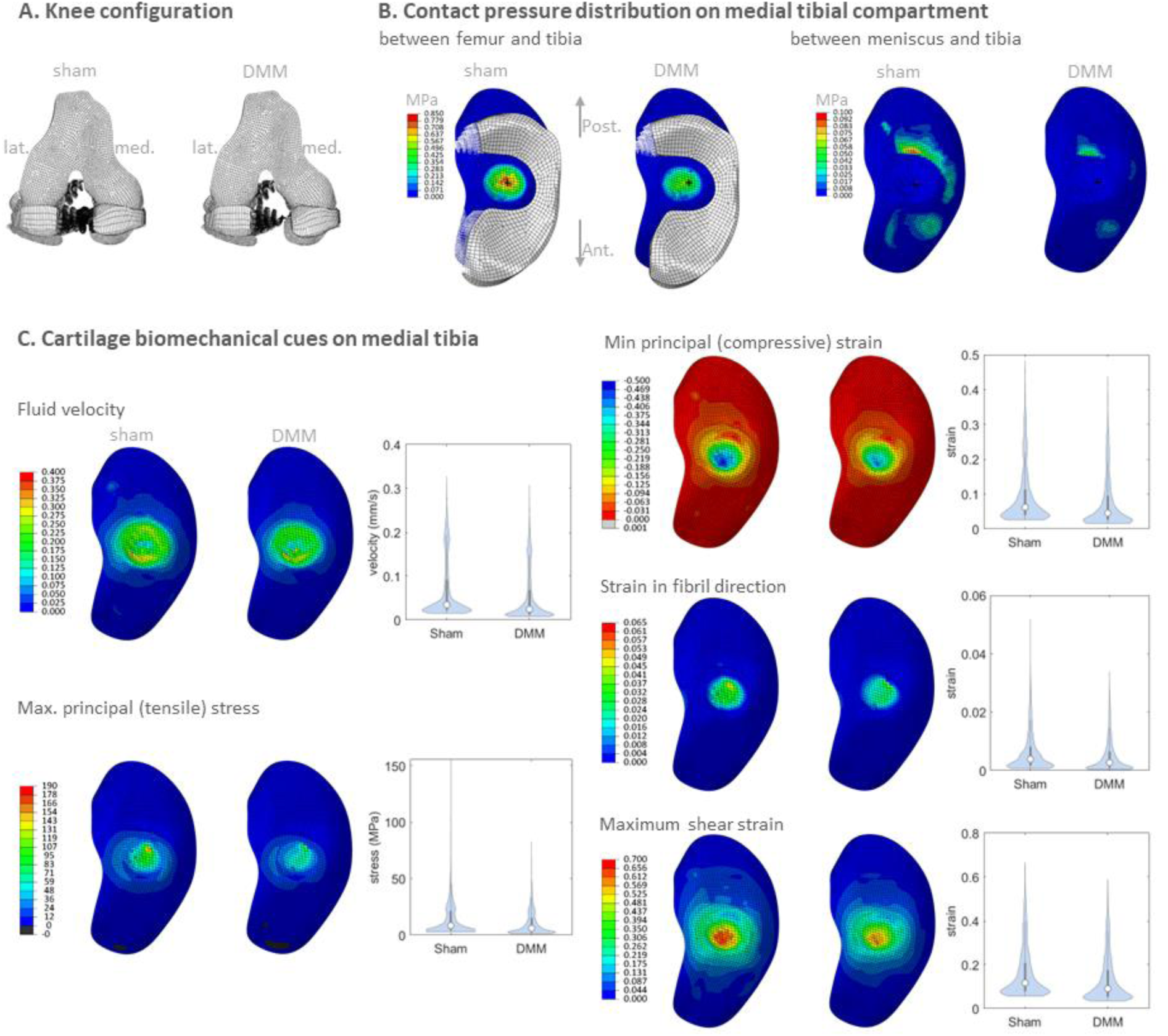
FE model analysis. **(A)** Front view of knee configuration at peak knee contact force in sham surgery and DMM (where the attachment of the medial meniscus is deactivated). **(B)** Contact pressure between the medial tibial and femoral cartilages, between the medial tibial cartilage and medial meniscus, and position of the center of pressure (black “+” sign). (**C**) spatial estimation and violin plots of mechanical cues associated with cartilage degeneration. For each mechanical cue, the upper half of nodes with the highest loading are drawn on the violin plots, to focus on the representation of the loaded region. Within this population, medians are plotted with a white dot and quartiles with a grey bar. The lower half of the nodes with the least loading are excluded from the violin plots.

## Discussion

Mechanical loading is widely accepted, but still poorly understood as a driver of osteoarthritis onset and progression. DMM is the current gold-standard rodent model of post- traumatic osteoarthritis, which is often described as a model of knee instability with locally increased loading. Yet, multiscale joint and tissue-level knee mechanics following DMM remain to be characterized, given the lack of high-quality, species-specific gait data. Our study addressed this gap by developing dedicated experimental setups and modeling methodologies to evaluate the multiscale mechanical environment in the knee joint of rats with DMM and sham surgery. Using experimental gait data and musculoskeletal modeling, we estimated knee poses and joint-level loading. We then used these data as inputs to a custom-made FE model, to spatially estimate tissue- level cartilage mechanical response. Although instability was observed in the frontal plane, joint- level loading and tissue-level mechanical cues related to cartilage degeneration did not increase but rather decreased with DMM compared to sham surgery.

Our study confirms frontal plane instability during gait following DMM, as documented by the knee adduction angle. Indeed, thanks to the use of biplanar fluoroscopy and implanted bone markers, combined with the musculoskeletal modeling workflow customized to account for 3D rotational motions, our study reports joint poses during gait that are more accurate compared to previous studies relying on skin markers ^23^.

This study is the first to document that joint-level knee loading is not increased in rats following DMM. Based on the comparable magnitude of the knee contact force, contact force on the medial compartment, and impulse on the medial compartment between both groups, we could not confirm increased joint level loading following DMM. This is because the associated magnitude of ground reaction forces and muscle forces used to control the knee joint during gait were similar.

Multiscale analysis of the spatial mechanical cues in the knee cartilage did not indicate increased tissue loading following DMM. DMM resulted in lower contact pressure on the tibia cartilage compared to sham surgery but with an identical location of the center of pressure, indicating no shift in the load-bearing area – a hypothesis often formulated to explain cartilage degeneration in the presence of joint instability. Furthermore, DMM reduced the biomechanical cues typically associated with cartilage degeneration. This may be caused by the different kinematic and kinetic profiles used as inputs to the FE model, associated with the representative gait profile of rats with DMM (in particular, lower resultant adduction moment).

The observed osteoarthritis-like structural changes in cartilage and subchondral bone are therefore not associated with alterations in the spatial distribution of the multi-scale mechanical loading environment and excessive mechanical joint and tissue-level loads could not be confirmed. Therefore, the degenerative changes observed in DMM may be explained through the combined effect of decreased mechanical loading and eventual inflammatory changes: the underloading observed at the tissue level in DMM suggests unloading of the cartilage tissue with insufficient mechanical cues in the cell microenvironment contributing to impaired matrix synthesis. Future research combining spatial-omics techniques with the spatial mechanical environment as evidenced by the insights of our model, has the unique potential to elucidate the unique role of the mechanical loads and molecular changes in the cells in directing the cartilage mechanome towards driving the onset and progression of OA.

Despite our efforts to validate the comprehensive multiscale modeling workflow, the results might still be subject to modeling-related limitations. Regarding the musculoskeletal model, validation of joint contact forces against experimental measures is not feasible, as this would have been too invasive to allow natural gait; however, the estimated values are consistent with what has previously been reported in the literature. The loading conditions used as input for the FE model - more specifically the normalized axial force - were considerably higher than the axial loads used in previous models of the mouse and rat knees, which were not based on species-specific musculoskeletal analysis (2.44 and 0.45 body weight, respectively) ^18,20^. Therefore, we need to question the mechanical loading landscape previously reported in the literature. Our current results report differences in mechanical cues between DMM and sham surgery, assuming identical mechanical cartilage behavior between both conditions. As different sets of material parameters may alter this comparison, our multi-scale modeling workflow provides a platform for future studies on the impact of disease-stage-specific mechanical cartilage behavior on the spatial relationship between cartilage loading and mechano-adaptive changes.

To conclude, using high-fidelity experimental data in combination with a dedicated multi- scale modeling workflow, we could not confirm that increased joint- and tissue-level knee loads drive the observed knee cartilage and bone degradation. The model-based findings on multiscale cartilage loading highlight the need to further study the interaction between mechanical and biological changes in the joint, to fully understand the complex mechanisms underlying cartilage degeneration in the onset and progression of osteoarthritis. Understanding these mechanisms may ultimately lead to the development of more effective prevention and treatment strategies for this debilitating condition. To catalyze this endeavor, we openly share experimental gait data and the modeling workflow, including musculoskeletal models and FE meshes.

## Acknowledgments

We appreciate the help of our colleagues at the KU Leuven: Peter Smolders (design of the walkway replica), Jente Van Havermaet and Peter Holland (manual labeling of fluoroscopic images), Natalia Ferreras-Moreno (animal care and behavioral training), Lies Storms (histology), Prof. Greet Kerckhofs (hexabrix scans), Dr. Frederique Cornelis (advice on animal experiments). We thank Jan Scholliers from University of Antwerp for designing and constructing the walkway. We are grateful to Prof. Nicolai Konow form University of Massachusetts Lowell for training on radio-opaque bead implantation.

## Author contributions

Conceptualization: JP, IJ

Methodology: JP, AE, SV, FM, GO, PA, SVW, IJ

Formal analysis: JP, FM, IJ

Investigation: JP, AE, FM, MVN, SVW

Resources: RL, PA, SVW, IJ

Visualization: JP, FM

Supervision: JP, RL, PA, SVW, IJ

Writing—original draft: JP, IJ

Writing—review & editing: JP, SV, FM, GO, MVN, AE, RK, PA, SVW, IJ, RL

Funding acquisition: JP, IJ, PA, SVW, RL

Final approval of the article: JP, SV, FM, GO, MVN, AE, RK, PA, SVW, IJ, RL

## Data and materials availability

Data and models have been uploaded to the KU Leuven Research Data Repository (RDR): https://doi.org/10.48804/2RTAOJ

