## Supplementary Materials for "Joint and tissue mechanics in post-traumatic osteoarthritis: insights from the rat model"

### Title

### Running title

multiscale knee biomechanics after DMM

### Authors

Judith Piet [1,2], Amir Esrafilian [3], Seyed Ali Elahi[1,4], Falk Mielke [5], Maarten Van Nuffel [2,6], Gustavo Orozco [3], Sanne Vancleef [1], Rik Lories [2,7], Rami K. Korhonen [3], Peter Aerts [5], Sam Van Wassenbergh [5], Ilse Jonkers [1,2]*

**Affiliations**

1 Department of Movement Science, KU Leuven, Belgium

2 Department of Development and Regeneration, Skeletal Biology and Engineering Research Centre, KU Leuven, Belgium

3 Department of Technical Physics, University of Eastern Finland, Finland

4 Department of Mechanical Engineering, KU Leuven, Belgium

5 Department of Biology, University of Antwerp, Belgium

6 Department of Orthopedic Surgery, University Hospitals Leuven, Belgium

7 Department of Rheumatology, University Hospitals Leuven, Belgium

### Surgeries

*Study design*

The sample size was based on a comparable study by Britzman et al., which evaluated knee biomechanics (in particular the knee contact force on the tibial medial plateau) in rats with tibial osteotomy and sham surgery, using reflective markers glued onto the skin and musculoskeletal modeling (n=5 rats per group) [1]. We added one additional rat per group to account for possible sample attrition (total of n=6 rats per group).

Our approach for deciding the sequence of surgeries was influenced by the availability of the micro-surgeon. While not completely randomized, we ensured that sham surgery and DMM were not systematically performed in a specific order. This was done to minimize potential confounders associated with the sequence of treatments. Regarding behavioral training, we randomized (simple randomization) the order in which the rats were trained every day over two weeks. The first author was aware of the group allocation during the conduct of the experiment and data analysis.

*Preparation*

Rats were anesthetized with isoflurane (5% at induction, then 2%), and kept on a warm pad. Eye lubricant was applied to prevent the eyes from drying, and analgesics were injected subcutaneously (Buprenorphine, 0.05mg/kg). Rats were draped, and the surgical site was shaved and disinfected with 70% ethanol.

*Destabilization of the Medial Meniscus or sham surgery*

The patellar tendon was identified visually after ethanol application, and manually by palpation. The skin and patellar tendon were incised with a #15 blade towards the medial side, on a length of a few millimeters, and spread open with dissection scissors. Bleeding was reduced by applying pressure with gauze, and recalcitrant bleeding was stopped with a drop of epinephrine (1:1000). In the group with DMM, the medial menisco-tibial ligament was transected (blade #15), as well as the anterior-medial attachment of the medial meniscus to the tibial plateau by a few millimeters. After the replacement of the extensor muscles, one drop of isobetadine was applied to the incision for sterility. The patellar incision was sutured with resorbable coated Vicryl sutures, and the skin incision was sutured with Ethilon polyamide sutures. Painkillers were administered for 48 hours post-surgery (Buprenorphine, 0.05 mg/kg, SC injection).

*Implantation of radio-opaque markers*

The lateral side of the thigh was palpated to identify the mid-part of the femur. A small skin incision was performed (less than 5 mm) and spread open to expose the biceps femoris. A small incision (a couple of mm) was performed in the direction of the muscle fibers and spread open with dissection scissors to expose the underlying femur. Pressure was applied with gauze to stop bleeding. A small manual bone drill was used to drill holes in the femur cortex, and 3 to 5 lead-free tin beads (0.5 mm diameter) were pushed into these holes with thin forceps. One drop of isobetadine was applied to the incision for sterility. The muscle incision was sutured with resorbable coated Vicryl sutures, and the skin incision was sutured with Ethilon polyamide sutures. The medial side of the tibia was palpated to identify the location where the tibia is closest to the skin. A small skin incision was performed (less than 5 mm) and spread open to expose the subcutaneous medial surface of the tibia. A small incision (a couple of mm) was performed in the direction of soft tissue covering the bone and spread open with dissection scissors and ease the drilling of holes. 3 to 5 beads were implanted, and the incision sites were sutured as described above. Painkillers were administered for 48 hours post-surgery (Buprenorphine, 0.05 mg/kg, SC injection).

### Ground reaction forces and biplanar fluoroscopy acquisition

Force plate outputs were amplified and acquired with a data acquisition system (DAQ). The force plate was calibrated before the measurements [2]. The space was calibrated with a calibration block, and X-ray image distortion was corrected with undistortion grids [3]. The coordinates of the force plate were identified using radio-opaque beads mounted on Lego bricks, and the direction of gravity was identified with a radio-opaque thread on which a weight was hanging. In one day, the gait of 2 to 3 rats was recorded. Being in the walkway with another rat was found to be a stronger incentive than the food rewards used during the behavioral training.

### Sample imaging and histology

At the end of each recording day, rats were sacrificed by overdose of Phenytoin/pentobarbital (Euthasol) followed by cervical dislocation. The right hindlimb was dissected into one intact piece, from the foot to the pelvis. Hindlimbs were fixed at an angle of 120 degrees, in 4% formaldehyde at room temperature for 5 days, with daily solution change. Full hindlimbs were imaged with a coarse resolution (GE Nanotom M, 40 micrometer voxel size, 90 kV, 200 μA). Samples were then incubated for one week in 40% ioxaglic acid (Hexabrix) solution. Knee structures were then imaged with a fine resolution (GE Nanotom M, 6.5 micrometer voxel size, 60 kV, 350 μA). Knees were then decalcified using 4% formic acid for two weeks (with daily changes on weekdays), embedded in paraffin, and sectioned for histological evaluation.

### Musculoskeletal modeling

*Degrees of freedom of the knee*

The generic model of Johnson et al. had one degree of freedom at the knee, with simmspline functions to define 3 translations and the 2 secondary rotations. The generic model was adapted in OpenSim 4.4. The knee center was defined as the midpoint of the long axis of the cylinder that fits through the femur condyles. Anatomical child and parent frames for the knee joint were created by offsetting frames relative to the femur and tibia frames (translation to the knee center for the tibia, translation, and orientation for the femur so that the mediolateral axis is aligned with the long axis of the cylinder that fits through the femur condyles and the longitudinal axis is aligned with the long axis of the femur). This generic musculoskeletal model of Johnson et al. was developed with female rats that were lighter than the male rats in the current study. Thus, the maximum isometric force of the muscles was tripled in the adapted musculoskeletal model. The secondary degrees of freedom (adduction and rotation) of the ankle were locked because bone landmarks were harder to identify on the foot, due to the overlap of the foot with the floor on X-ray images.

*Static optimization*

Muscles were modeled as ideal force generators, thereby ignoring the force-length and force-velocity characteristics, because the parameters describing the force-length velocity-relationship in rat muscles are largely unknown. Muscle force distribution was determined so that the muscle moments balance the joint torques (inverse dynamics) while minimizing the sum of squared muscle activations. Because knee secondary degrees of freedom (adduction and rotation) are also stabilized by bone-on-bone contact and knee ligaments, large reserve actuators were used for the secondary degrees of freedom. The knee flexion reserves reported in this study were negligible compared to the external knee flexion moment, which indicates that knee flexion is indeed actuated by muscle forces.

*Validation*

For the ground reaction forces, the maximum vertical ground reaction forces for rats with sham surgery ranged from 82% to 113% body weight. This is in line with, although slightly higher than values previously reported in the literature, with approximate average values of 60% body weight in male Wistar rats [4], 70% in Long-Evans rats [5], 80% in male Sprague-Dawley rats [1]. Ground reaction forces may be higher because rats in our study ran faster (50-95cm/s in this study, versus 30-50cm/s in [5]). For the knee kinematics, the peak knee flexion angle for rats with sham surgery ranged from 77 to 97 degrees, which is similar to what has previously been reported in the literature (on average 96.4 degrees at 30cm/s, 99.4 degrees at 60 cm/s [6]).

The order of magnitude for flexor moments is the same as previously reported values (-15 to -2 BW.mm on average in female Long-Evans rats [7]). The knee adduction moment rats with sham surgery ranged from -43 to 33 BW.mm, while Britzman et al. report a range of -1 to 5 BW.mm on average [1]. The estimated mean magnitude of the knee contact force of 14 BW for rats with sham surgery is in the same order of magnitude as what has been reported by Wehner et al. (7.5 BW) [8]. Wehner et al. reported a lower magnitude of the knee contact force, which is consistent with the lower ground reaction forces used in their study [5]. The muscle forces estimated by the static optimization step are in the same order of magnitude as values reported in previous studies. The estimated mean peak forces for the rectus femoris, vastus lateralis, lateral gastrocnemius, and medial gastrocnemius in rats with sham surgery are 3.7 BW, 5.4 BW, 3.8 BW, and 1.6 BW respectively. Values reported in literature estimated by musculoskeletal modeling [8] and *in vivo* measurements [9] are approximately 7 BW, 4 BW, 3 BW, and 1.3 BW respectively.

### FE model

*Meshes and assembly*

Knee cartilages, menisci, and ligament insertion points were manually segmented (Mimics, version 21, Materialise, Belgium) using the fine-resolution hexabrix micro-CT scan of a sham knee, post-processed (3-Matic, version 16, Materialise, Belgium), and the 3D geometries were meshed using hexahedral elements in Ansa (v21.0.1, BETA-CAE Systems, USA) and HyperMesh (v2019, HyperWorks, USA). Bones were assumed rigid compared to cartilages and excluded from the model. All the meshed geometries were then imported into the Abaqus software (version 21, Dassault Systemes, USA) to create a complete FE model of the rat knee. Articular cartilage and menisci were modeled with porous 8-node elements (C3D8P), and mesh density was set according to a previously published model of the rat knee [10]. All the parts were assembled, and ligament bundles and menisci horn attachments were defined according to the insertion points obtained from the segmentation. The reference points, node, and element sets required for applying boundary conditions, loads, and contacts were defined. The FE model reference point coordinate system aligned with the musculoskeletal model **(Fig. 4B)**. Interactions (general contact) and couplings were defined. Reserve stiffnesses were defined to account for anatomical structures not included in the model [11].

*Material properties*

Ligaments and menisci horn attachments were modeled as bi-linear spring bundles [12]. Femoral and tibial cartilage was modeled with fibril-reinforced poro-viscoelastic properties, and menisci were modeled with fibril-reinforced poroelastic properties. The primary collagen fibrils in the cartilage were arranged according to the depth-dependent Benninghoff arcade pattern. The menisci fibril network aligned circumferentially. In both tissues, fluid flow through permeability was assumed to be strain- and void ratio-dependent. Cartilage and menisci were modeled using the equations detailed in [13,14]:

- Young's modulus of the non-fibrillar matrix: 4.2 MPa for cartilage, 0.5 MPa for menisci [15]
- Poisson's ratio of the non-fibrillar matrix: 0.42 for cartilage [16], 0.36 for menisci [15]
- Initial modulus of the fibril network: 16.17 MPa for cartilage, 28 MPa for menisci [15]
- Strain-dependent modulus of the fibril network: 150 for cartilage [17]
- Damping coefficient of the fibril network: 1062 for cartilage [13]
- Initial permeability: $3.3 \times 10^{-15}$ $m^4/Ns$ for cartilage [10], $1.2 \times 10^{-15}$ $m^4/Ns$ for menisci [18]
- Ratio between the primary and secondary fibrils: 12.16 for cartilage and menisci [13]
- Permeability strain-dependency factor: 1.67 for cartilage [10], 12.16 for menisci [19]

*Loading and boundary conditions*

The FE model's reference point was the origin of the coordinate systems in the associated musculoskeletal model. The musculoskeletal and FE models coordinate systems were similarly defined to ensure the consistency of the kinematics and kinetics **(Fig. 4B)**. The bottom of the tibia was fixed. All the nodes located on the femoral cartilage to the subchondral bone interface were coupled to the femoral reference point. After bringing the parts in contact (step-1), knee flexion was applied to correct for knee over-extension in the micro-CT scan (step-2). The knee flexion angle and the resultant adduction moment and joint contact forces (axial, anteroposterior, and mediolateral) from muscles, inertial, and gravitational forces were then applied to the femoral reference point (step-3). The internal/external rotational degree of freedom was free and the rotation moment from muscles, inertial and gravitational forces was not applied, as it resulted in excessive knee rotation, as observed in previous knee models [20].

### Supplementary figures

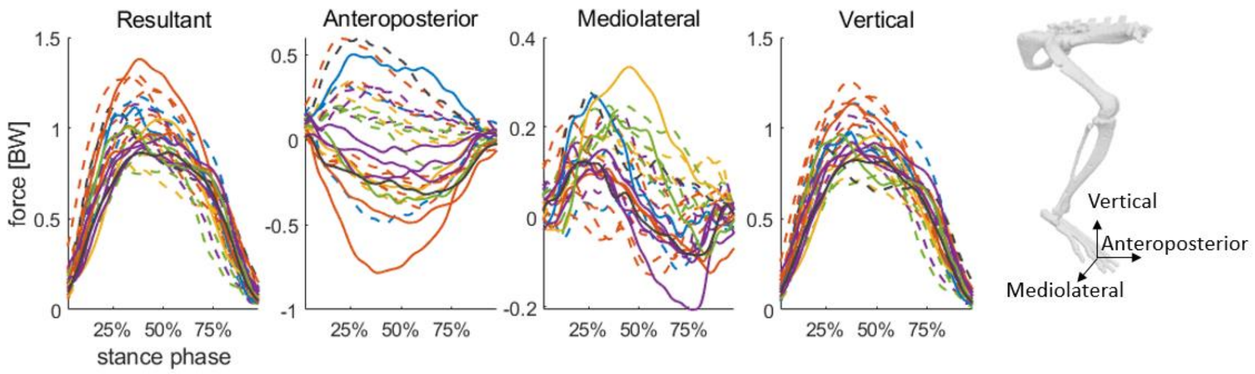


**Fig S1. Normalized ground reactions forces over stance phase**. N=6 rats, n=18 trials for sham; N=6, n=9 for DMM. Trials from rats with sham surgery are represented with a continuous line, trials from rats with DMM are represented with a dashed line. Trials from the same rat are represented with the same color.

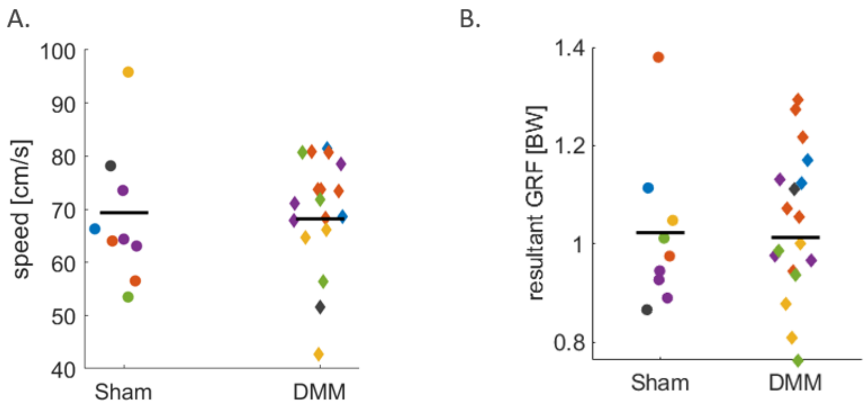


**Fig S2. Speed and normalized ground reactions forces.** No difference was observed in **(A)** ambulating speed and **(B)** normalized peak resultant ground reaction force (GRF) between rats with DMM and sham surgery. N=6 rats, n=18 trials for sham; N=6, n=9 for DMM. Trials from the same rat are represented by the same color. The mean for each group is represented with the horizontal black line.

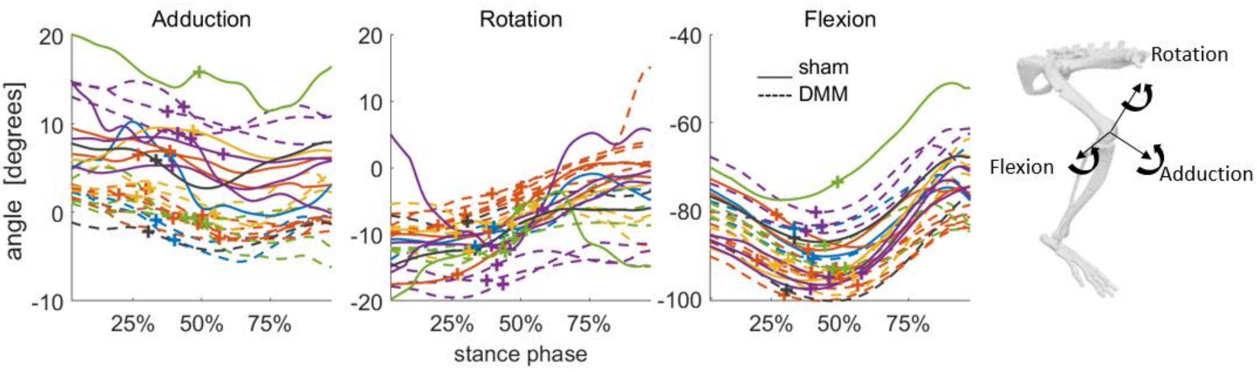


**Fig S3. Knee kinematics over stance phase**, obtained by inverse kinematics. N=6 rats, n=18 trials for sham; N=6, n=9 for DMM. Trials from rats with sham surgery are represented with a continuous line, trials from rats with DMM are represented with a dashed line. Trials from the same rat are represented with the same color. The data points marked with "+" correspond to the kinematics values when resultant normalized contact force is at its maximum.

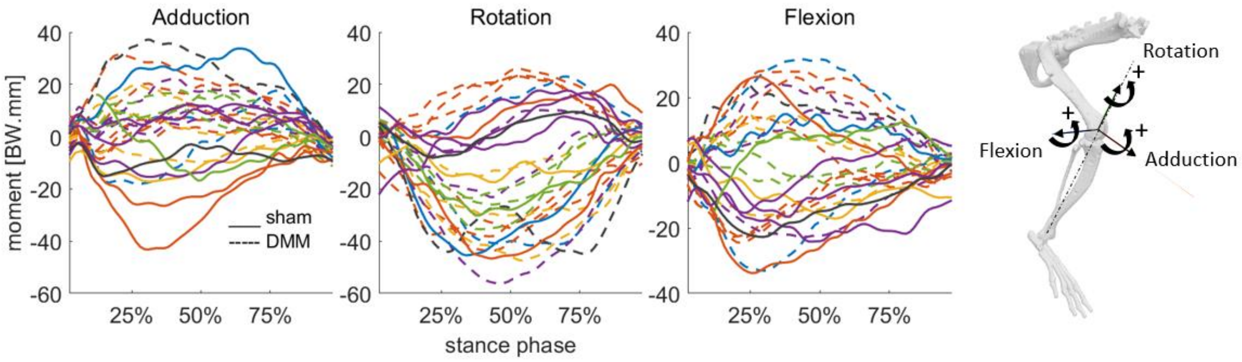


**Fig S4. Normalized external knee moments over stance phase**, obtained by inverse dynamics. N=6 rats, n=18 trials for sham; N=6, n=9 for DMM. Trials from rats with sham surgery are represented with a continuous line, trials from rats with DMM are represented with a dashed line. Trials from the same rat are represented with the same color.

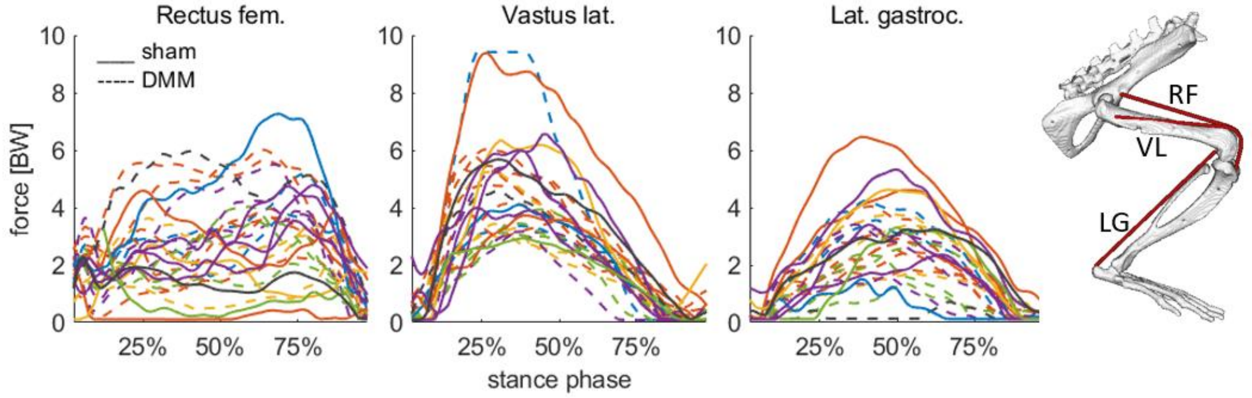


**Fig S5. Normalized forces for muscles spanning across the knee**, obtained by static optimization of muscle forces. N=6 rats, n=18 trials for sham; N=6, n=9 for DMM. Trials from rats with sham surgery are represented with a continuous line, trials from rats with DMM are represented with a dashed line. Trials from the same rat are represented with the same color.

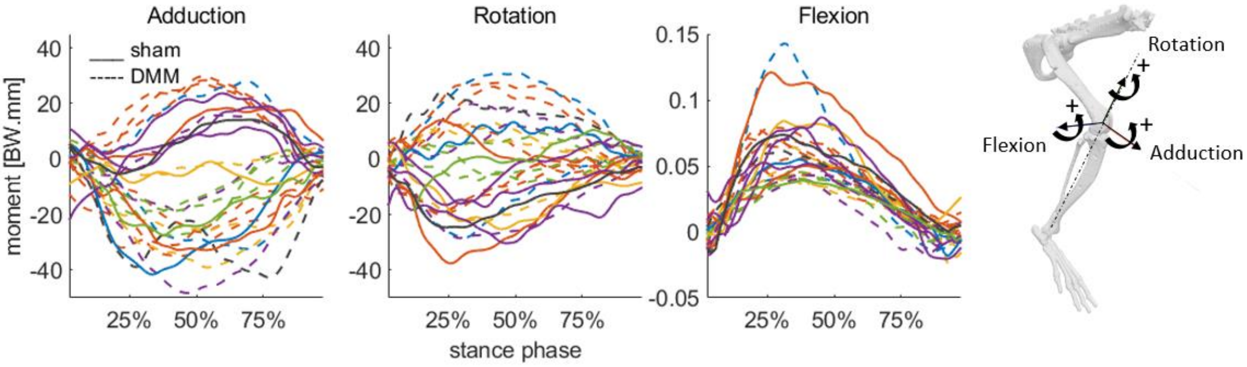


**Fig S6. Normalized reserves torques**, using during static optimization of muscle forces. N=6 rats, n=18 trials for sham; N=6, n=9 for DMM. Trials from rats with sham surgery are represented with a continuous line, trials from rats with DMM are represented with a dashed line. Trials from the same rat are represented with the same color.

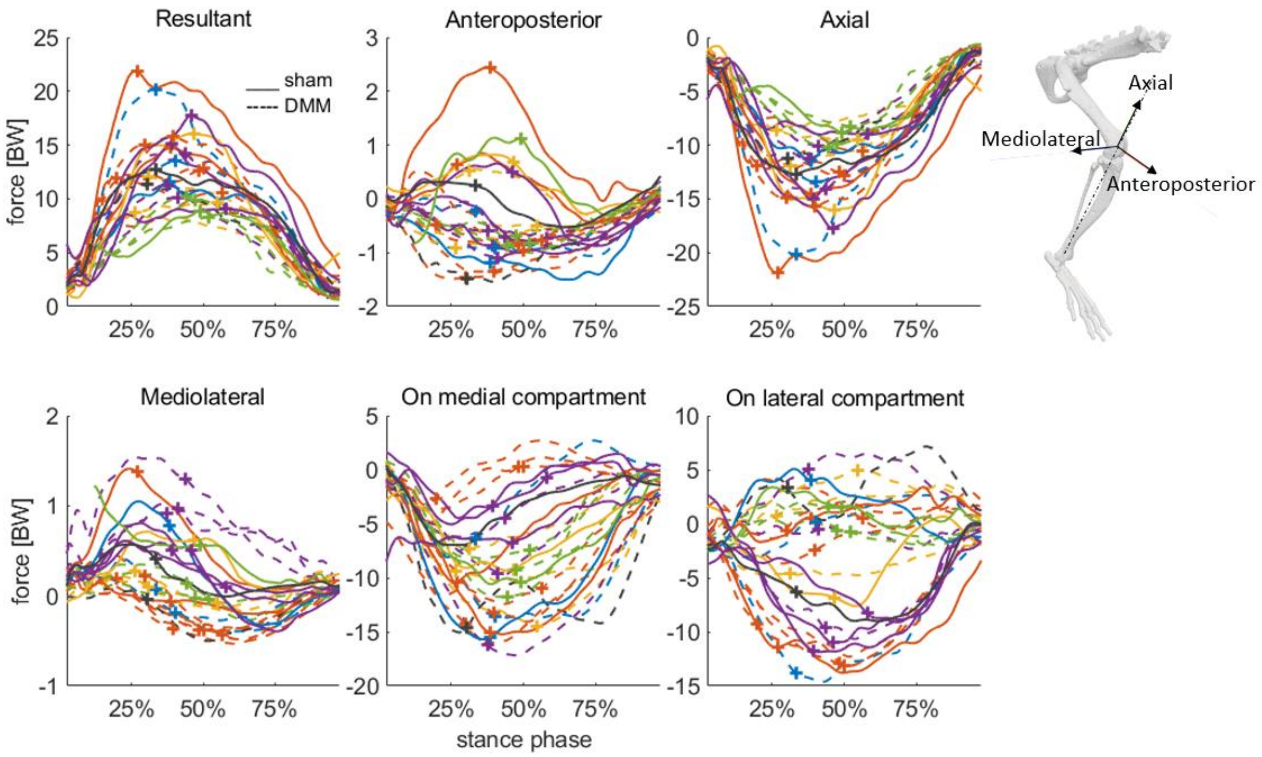


**Fig. S7. Normalized knee contact forces over stance phase**, obtained by joint reaction analysis. N=6 rats, n=18 trials for sham; N=6, n=9 for DMM. Trials from rats with sham surgery are represented with a continuous line, trials from rats with DMM are represented with a dashed line. Trials from the same rat are represented with the same color. The data points marked with "+" correspond to the kinematics values when resultant normalized contact force is at its maximum.
